# Evaluating Temporal Orders for Local Non-Stationary Biological Signals Analysis: A Python Framework and Simulation Study

**DOI:** 10.64898/2026.07.26.740785

**Authors:** Marcel Młyńczak, Maciej Rosoł, Kacper Korzeniewski, Jakub S. Gąsior

## Abstract

**Background and Objective:** Accurately parameterizing dynamic, time-varying interactions in physiological systems is a methodological challenge, as global causal discovery methods may obscure transient, local fluctuations. This study introduces *tempord*, an open-source Python library designed to estimate local temporal orders and evaluate the short-term stability, directionality, and strength of causal links in non-stationary biological signals.

**Methods:** The algorithm estimates temporal relationships by keeping one signal stationary while iteratively shifting another one within a sliding window. To parameterize optimal inter-signal shifts (causal vector, CV), the framework utilizes linear modeling or time series distance metrics. The methodology was validated through a simulation study on synthetic bivariate signals with mathematically imposed dynamic phase delays, under both deterministic and noisy conditions. Furthermore, in-vivo capabilities were demonstrated by evaluating cardiorespiratory coupling dynamics across spontaneous and music-induced relaxation breathing states.

**Results:** The simulation study demonstrated that the extracted CV trajectories precisely aligned with ground-truth temporal delays, assessed using mean absolute error and root mean square error for both noise-free and noisy synthetic data. In-vivo application demonstrated dynamic temporal stability and the detection of minor step changes during autonomic nervous system state transitions.

**Conclusions:** The *tempord* Python package bridges the gap between global causal discovery and local beat-by-beat statistical parameterization. It provides a robust “bottom-up” analytical instrument for investigating the transient mechanisms governing complex biological networks.

## I. Introduction

THE human organism functions as an integrated network of physiological systems whose continuous, dynamic interactions underpin the maintenance of homeostasis [1], [2]. Accurately parameterizing these complex, time-varying interactions from observational data remains a significant methodological challenge. Standard approaches—including many machine learning (ML) models—focus predominantly on correlations or predictive accuracy, often failing to reveal underlying causal mechanisms [3]. Furthermore, traditional “global” causal discovery methods, such as Granger causality or transfer entropy, average out “local” fluctuations when applied over entire recording sessions. This aggregation obscures the transient nature of physiological couplings [4], [5].

A fundamental limitation of many dynamic causal models is the assumption of an *a priori* known causal network structure. While feasible at a macroscopic level, this requirement is rarely satisfied when analyzing evolving biological systems directly at the signal level [6]. To overcome the limitations of static structural assumptions, modern computational frameworks, such as the Temporal Causal Discovery Framework (TCDF) [3] and Dynamic Causal Network Autoregression (DCNAR) [6], have begun integrating dynamic causal inference.

To address the limitations of static structural assumptions, modern computational approaches have begun integrating dynamic causal inference. For instance, the Temporal Causal Discovery Framework (TCDF) utilizes attention-based convolutional neural networks to discover causal relationships and delays directly from time series [3]. Similarly, Dynamic Causal Network Autoregression (DCNAR) provides a two-stage approach that learns causal structures from data to enable time-varying causal inference and impulse response analysis without pre-specified networks [6]. Python frameworks for causal discovery were also proposed [7], as well as Granger-based approaches utilizing the non-linear recurrent deep learning techniques [8].

Building upon those paradigms and the need for “local” physiological assessment, this paper addresses the temporal orders estimation method [9], with the aim to introduce *tempord*, a new Python library designed to estimate local temporal orders in non-stationary biological signals [10]. Extending and refining our previous framework created for R language, we provide the analytical tool intended both for exploratory data analysis (mainly its visual outcome), as well as to evaluate the short-term stability, directionality, and strength of causal links within narrow time intervals.

The second objective of the study is to present the algorithm validation on synthetic signals with known phase delays to provide a structural ground truth. Subsequently, we also demonstrate the practical capabilities by evaluating cardiorespiratory coupling (CRC) dynamics illustrating how the approach utilizing temporal orders can bridge the gap between high-level causal discovery and the precise parameterization of local temporal stability.

## II. Methods

### A. The “tempord” algorithm

The core of the algorithm [9] is designed to estimate and evaluate (both quantitatively and visually) local, short-term so-called temporal orders (TO) between two physiological signals measured synchronously. Given a bivariate dataset, the fundamental mechanism involves keeping the first signal (a specific segment, e.g., 10 seconds) computationally stationary while the second signal is iteratively shifted backward and forward in time within a predefined, user-specified shift range, e.g., from -2 to 2 seconds and shift step. This range does not necessarily have to be symmetrical relative to the zero shift.

The analysis is performed within sliding signal windows of a fixed length. To quantify the temporal relationship at each shift, the algorithm supports two primary mathematical approaches:

- **Linear Modeling (LM):** Evaluates the relationship using the adjusted *R*^2^ measure to determine the optimal fit.
- **Time Series Distance (TD):** Computes proximities using metrics such as Euclidean, Manhattan, Chebyshev, Correlation, and Cosine distance measures.

For every discrete point in time, the algorithm computes a parameter array across the whole shift range. For the analysis, either the raw signal or its instantaneous phase, calculated via the Hilbert transform, can be selected.

To parameterize the optimal inter-signal shift, a so-called Causal Vector (CV) is extracted by defining the curve of maximum values (for the LM approach) or minimum values (for the TD approach) across the shift range provided.

To ensure analytical robustness and filter out insignificant noise, thresholding can be applied; then, only values strictly above the LM threshold or below the TD threshold are utilized to trace the final causal vector. Consequently, fragments where the CV remains non-intermittent and near-parallel to the X-axis signify periods of stable temporal coupling.

The method itself is designed to be applied on bivariate datasets; however, it can be easily extended to more signals within the one-versus-one estimation paradigm. The module is capable of performing this task when multiple columns are provided.

The analytical scheme of the algorithm is presented on Figure 1. The algorithm is also described by the pseudo-code in Listing 1.

**Listing 1.**
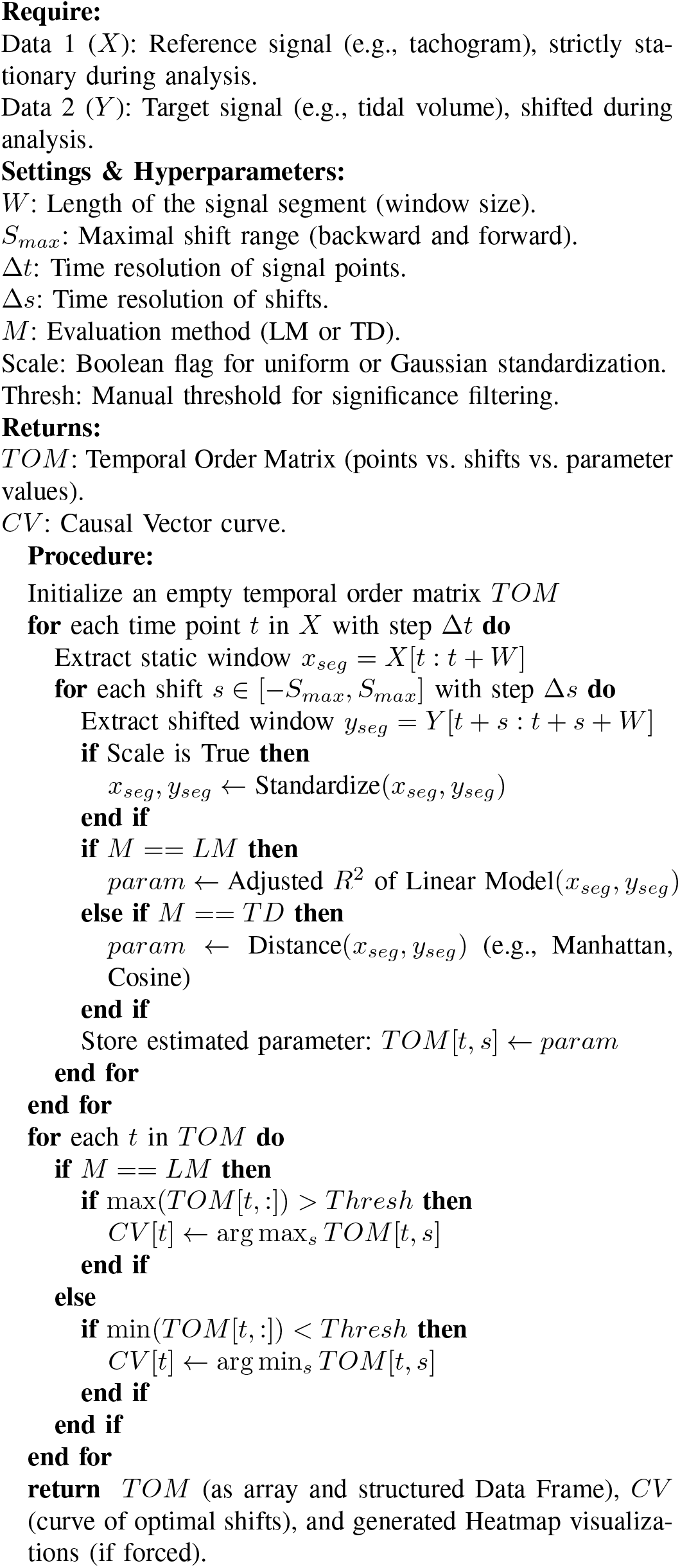
Local Temporal Orders and Causal Vector (CV) Estimation.

**Fig. 1.**
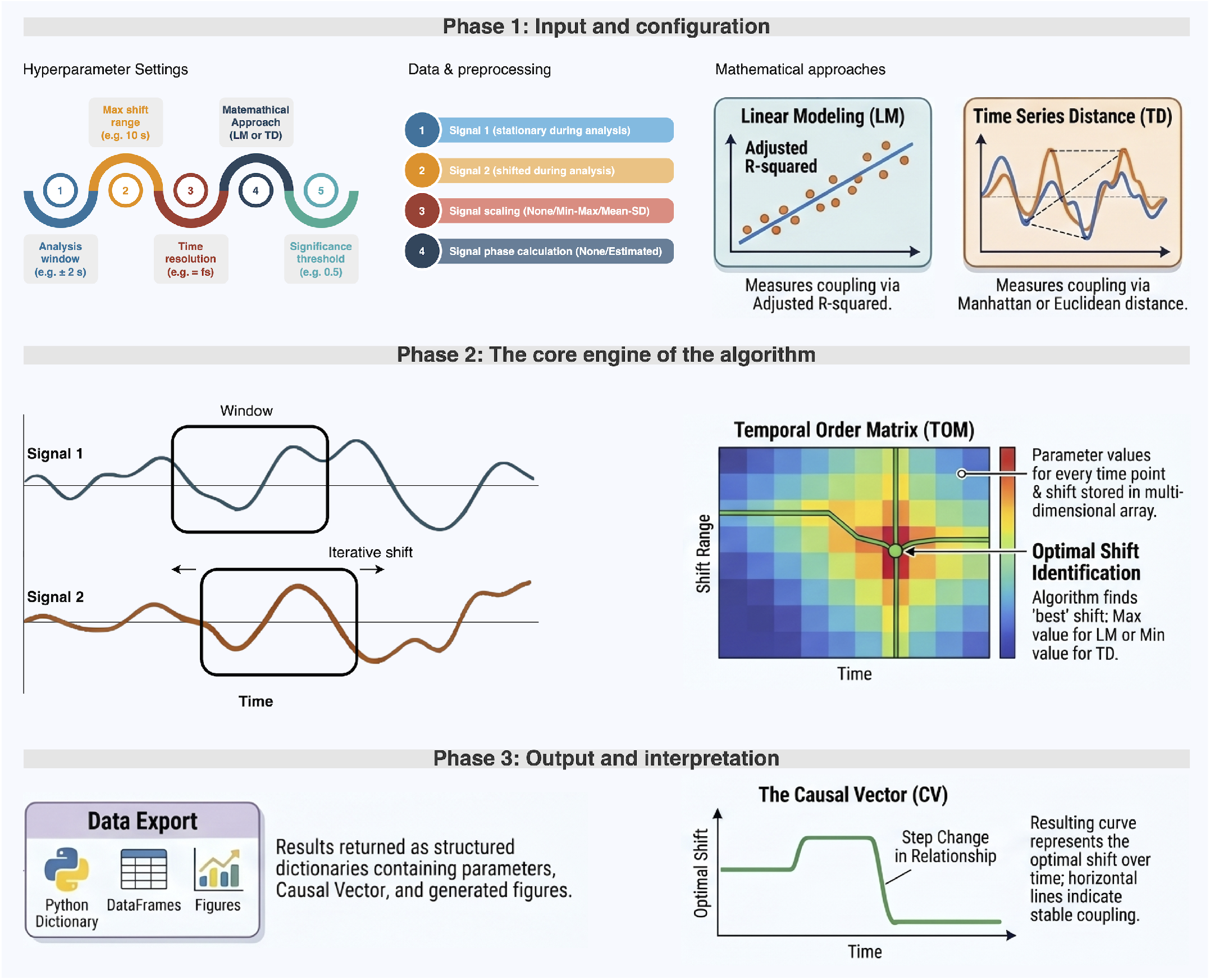
The illustration of the algorithm analytical scheme.

### B. Python Package Architecture

To overcome the limited adoption of the previous R-based implementation and facilitate integration into modern data science pipelines, the methodology was entirely re-architected as an open-source Python package (*tempord*) [10]. The library is publicly available via the Python Package Index (PyPI) and includes interactive Jupyter Notebook (comprising the examples demonstrated in the manuscript) to ensure a low barrier to entry for rapid adaptation [11].

The Python implementation leverages vectorized operations to handle the procedure efficiently. It is designed to operate on either raw physiological signals or their extracted phases (an approach similar to that utilized in the *neurokit2* Python package [12]). Furthermore, the package incorporates automated visualization modules to generate highly interpretable bivariate temporal order heatmaps and CV trajectories, enabling researchers to visually pinpoint regions of stable coupling and step changes caused by extrinsic study conditions.

The core `tempord` function accepts the following arguments:

- `df (pandas.DataFrame):` A data frame in which the columns represent consecutive signals to be analyzed. All signals should have the same length and sampling frequency. If the data frame contains more than two columns, the analysis is performed for each pair of signals.
- `method (str):` The mathematical approach used to calculate temporal orders.
- `modality (str):` The signal representation to be used. The available options are “raw” and “phase”.
- `scaling (int):` The scaling method applied to each analysis window. If set to 0, no scaling is applied. If set to 1, min–max normalization is used. If set to 2, z-score standardization is applied.
- `sig_length (float):` The length of the analyzed window, expressed in seconds.
- `max_shift_seconds (tuple):` A tuple containing two values that specify the maximum backward and forward shifts, respectively, expressed in seconds.
- `point_time_res (int):` The step size of the analyzed window, expressed in samples.
- `shift_time_res (int):` The step size of the shift between the values specified by `max_shift_seconds.`
- `make_figure (bool):` Indicates whether the results should be plotted.
- `td_type (str):` The type of time-distance metric to be used. This argument applies only when `method` is set to “TD”. The currently available options are “euclidean”, “manhattan”, “chebyshev”, “correlation”, and “cosine”.

The `tempord` function returns a dictionary in which the keys are tuples representing pairs of analyzed signals. These keys are constructed from the corresponding column names in the input df. Each value is a dictionary containing the following elements:

- `“Tempord” (pandas.DataFrame):` A matrix containing the calculated temporal-order parameter values.
- `“CV” (pandas.DataFrame):` Abbreviation for the Causal Vector; a data frame containing, for each analyzed time point, the shift values at which the maximum parameter value exceeded the predefined threshold.
- `“Fig” (matplotlib.figure.Figure or None):` A Matplotlib figure presenting the results if plotting is enabled; otherwise, None.

To allow users to quickly evaluate different thresholds without recalculating the entire temporal-order matrix, the *tempord* package provides a function called `get_causal_vector.` This function uses the temporal-order matrix, the selected method, and the threshold value as inputs and returns the corresponding causal vector. For convenience, the package also provides a `make_plot` function, which can be used to generate a plot of temporal orders with new CV values for a specified threshold.

### C. Simulation Study

To provide an objective, ground-truth validation of the algorithm, we designed a comprehensive simulation study. The objective was to strictly evaluate the algorithm’s precision in tracking dynamic phase delays before applying it to noisy biological data.

We generated synthetic signals *X*_1_(*t*) and *X*_2_(*t*) using a fundamental frequency *f* = 0.25 Hz, representing a generic physiological rhythm. The simulation was divided into three distinct temporal phases (*T*_1_, *T*_2_, and *T*_3_, each for 2 minutes) to model the dynamic, non-stationary nature of biological couplings, e.g., during different conditions. Across these phases, we introduced specific step changes in the time delay *τ* (*t*) between the signals, as well as slight, localized fluctuations in their respective amplitudes *A*_1_(*t*) and *A*_2_(*t*). We assumed the sampling frequency at the level of 25 Hz.

First, we constructed the noise-free deterministic signals governed by the following equations:

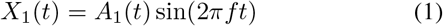

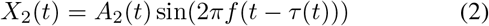

The dynamic time delay *τ* (*t*), representing the optimal shift, was mathematically imposed to shift from a positive lag to a negative lag across the three phases:

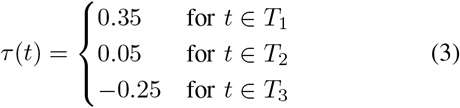

Concurrently, to ensure the algorithm’s robustness against amplitude modulation (also commonly occurring in physiological recordings), the amplitudes were programmed to vary slightly across the corresponding intervals:

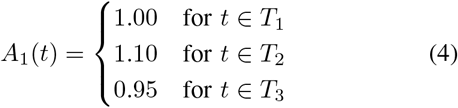

and

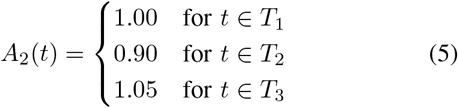

By establishing this noise-free baseline, we could confirm the algorithm’s fundamental mathematical accuracy in identifying the exact predefined temporal orders (350 ms, 50 ms, and -250 ms, respectively) without interference.

In the second stage of the simulation, we aimed to emulate the complexity and unpredictability of real-world biological recordings. To achieve this, independent zero-mean Gaussian noise components were injected into both generated time series. The resulting noisy signals, 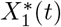 and 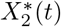, were defined as:

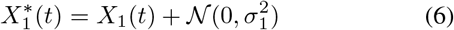

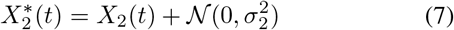

where 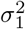 and 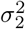 denote the variances of the noise, we assumed the values 0.08 and 0.12, respectively.

By possessing the exact mathematical ground truth, we could quantitatively evaluate the algorithm’s sensitivity and precision, directly comparing the trajectory of the estimated Causal Vector against the artificially imposed phase shifts.

To quantitatively evaluate the precision of the algorithm under both deterministic and noisy conditions, the deviation between the estimated Causal Vector (CV) and the true, mathematically imposed phase delay was rigorously measured. Specifically, the Mean Absolute Error (MAE) and the Root Mean Square Error (RMSE) were calculated across all simulated time points. These statistical metrics serve to validate the framework’s accuracy in tracking local temporal order transitions and to quantify its resilience to signal perturbations and stochastic noise.

### D. In-vivo Cardiorespiratory Study

Following synthetic validation, the framework was tested on in-vivo physiological data to demonstrate its capability in evaluating CRC [1], [2]. To ensure reproducibility and transparency, we utilized open-access data sourced from PhysioNet, comprising recordings under various measurement conditions to highlight physiological differences [13].

The analysis focused on two primary parameters: the continuous tidal volume (TV) curve, representing respiratory activity, and the tachogram (RR intervals), representing cardiac activity. We selected the Combined measurement of electrocardiogram (ECG), Breathing and Seismocardiograms Database (CEBSdb). This specific dataset was selected due to its rigorous, multi-phase protocol designed to analyze the autonomic nervous system. The protocol captures high-resolution physiological signals during both an initial baseline phase of spontaneous, unconstrained breathing and a subsequent relaxation phase induced by listening to classical music, which naturally lowers the respiratory rate and shifts autonomic dominance.

In order to simulate a continuous transition between physiological states, a unified time series was constructed by concatenating two distinct recording phases from the same subject. Specifically, we extracted equal five-minute segments from both the baseline phase (spontaneous breathing) and the subsequent classical music phase (induced relaxation). To prevent boundary artifacts of R-peak detections at the juncture, all signal preprocessing steps were executed independently on each segment. First, R-peaks were automatically extracted from the raw ECG signal to compute the consecutive RR intervals, yielding the tachogram, using *neurokit2* Python module [14]. Because the tachogram is inherently non-uniformly sampled in time, both the RR intervals and the continuous TV surrogate signal were resampled to a common, uniform temporal grid. We applied cubic spline interpolation to align both time series at a sampling frequency of 25 Hz. Moreover, to mitigate boundary artifacts introduced by the spline extrapolation, the initial and final two seconds of the interpolated data were systematically trimmed.

The fully processed segments were subsequently merged along a continuous temporal axis. This concatenated dataset enables the *tempord* framework to seamlessly evaluate the dynamic physiological adaptation, providing a continuous trajectory of the causal vector as the autonomic nervous system activity shifts from a standard resting state to a relaxation-induced regime.

Analyzing the shift from spontaneous to the music-induced relaxed state provides a robust in-vivo ground truth to empirically demonstrate how the algorithm detects step changes in CRC, confirming the relative precedence of the cardiac rhythm during slowed breathing rates.

For the algorithm settings, the analysis utilized a segment length of 10 seconds. The maximum temporal shift was constrained to 2 seconds in both forward and backward directions, a boundary defined by average human breathing frequencies and the known physiological characteristics of CRC phenomena [1], [2].

Moreover, we performed an exploratory analysis of the complexity and execution time, in the form of a graph of the length of time depending on the window length (from 1 to 10 seconds with 0.5 seconds step), sampling frequency (from 5 to 25 Hz with 5 Hz step), and range size (from -1;1 to -2;2 seconds range with 0.1 step). Each parameter configuration was evaluated three times, and the resulting execution times were averaged.

## III. Results

### A. Simulation Study Results

The signals utilized in simulation study are presented in Fig. 2.

**Fig. 2.**
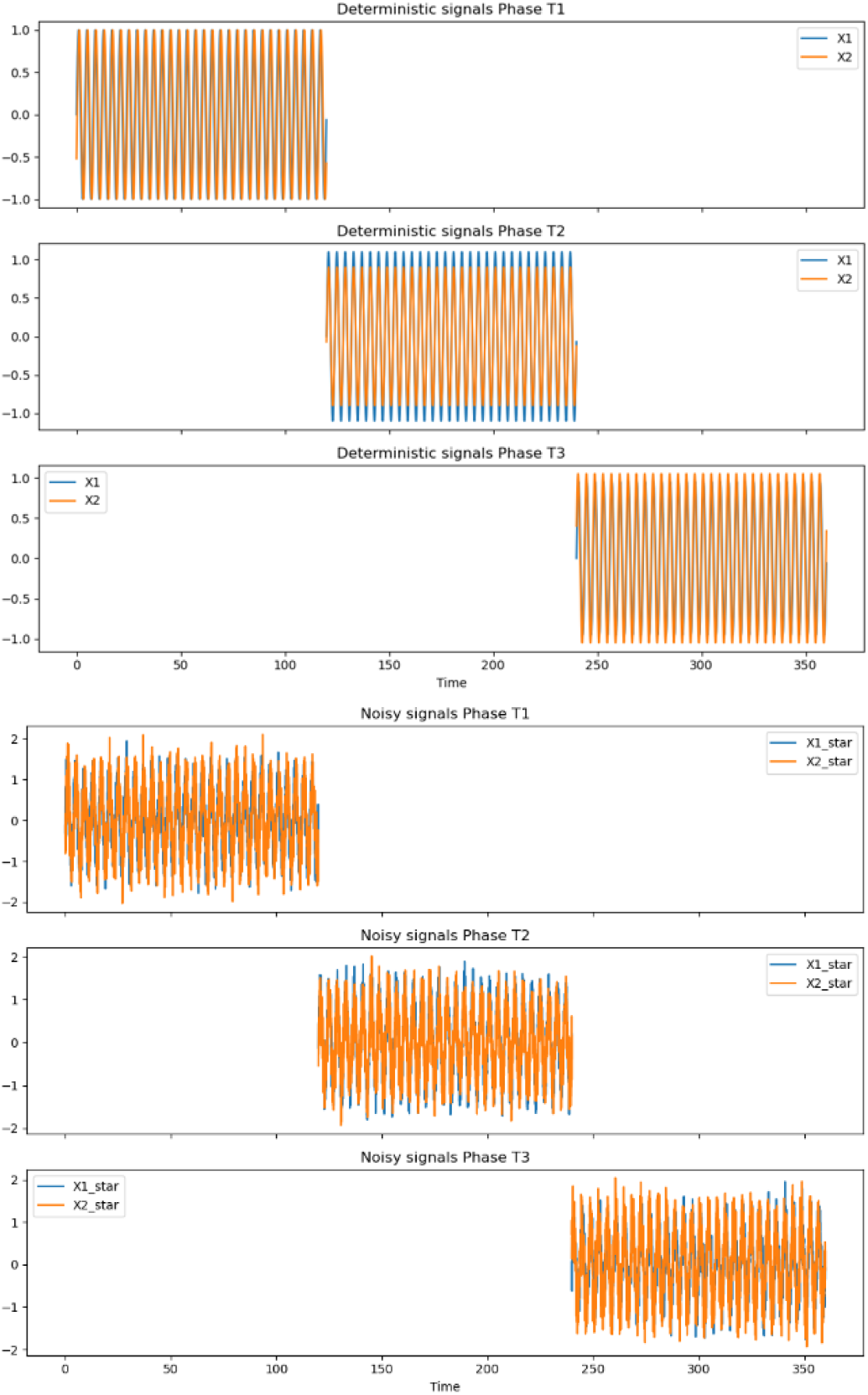
Simulated signals with different time shifts and amplitudes from three phases T1, T2, and T3 presented in the top, in the middle and at the bottom, respectively. Top: without noise; bottom: with noise.

The visualizations of bivariate matrices (as heatmaps) resulting from the *tempord* algorithm are presented in Fig. 3 and Fig. 4 for the pairs of *X*_1_(*t*) with *X*_2_(*t*) and 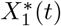 with 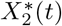, respectively.

**Fig. 3.**
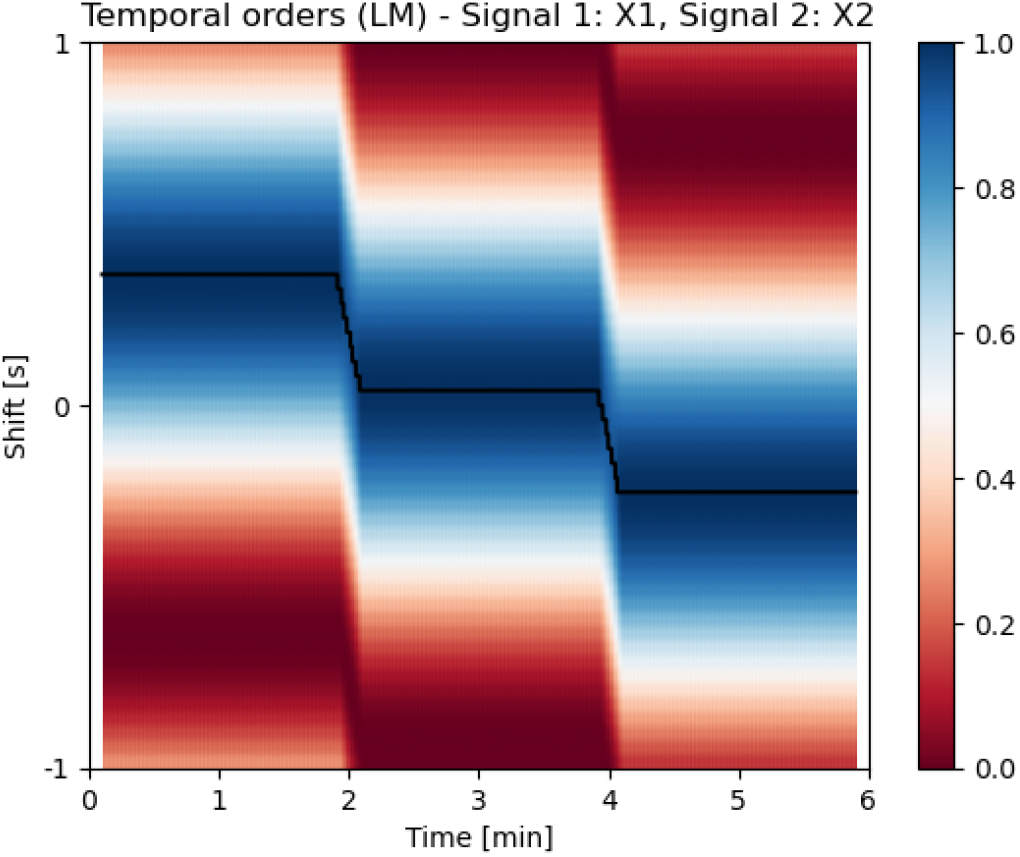
The temporal order heatmap for simulated signals without Gaussian noise added, with the CV extracted, as well as imposed phase delays.

**Fig. 4.**
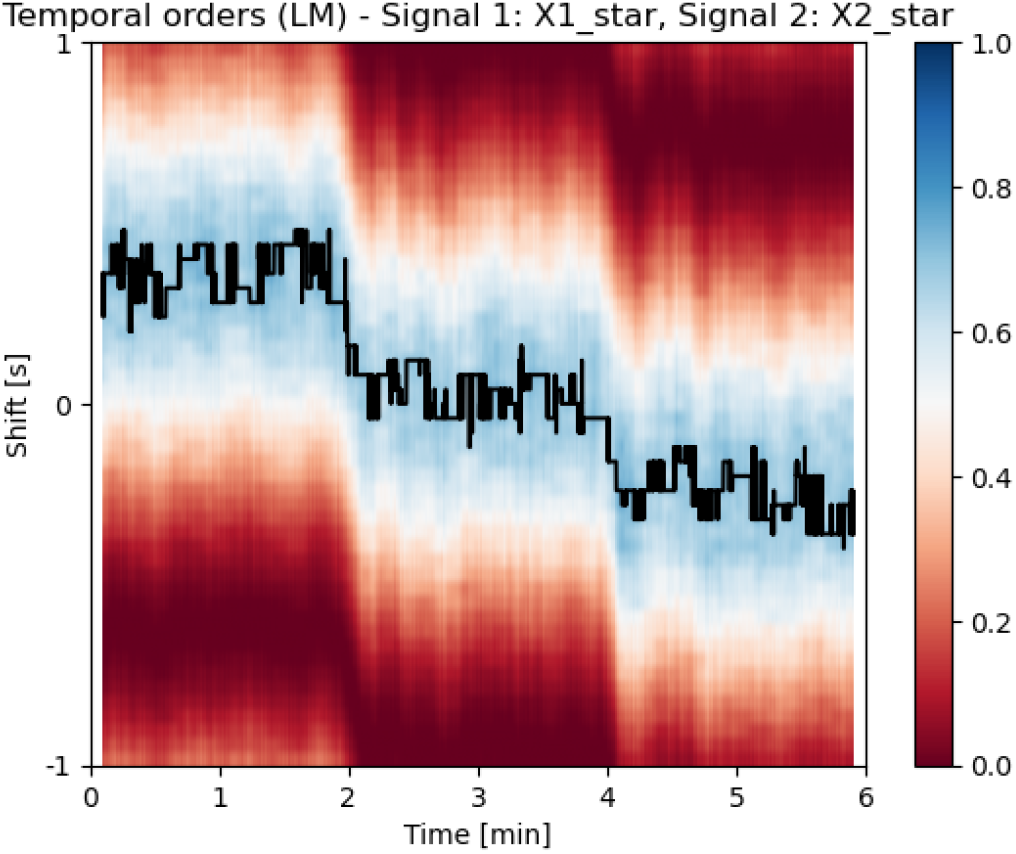
The temporal order heatmap for simulated signals with Gaussian noise added, with the CV extracted, as well as imposed phase delays.

The crucial outcomes of this simulation were the trajectory and values of the extracted Causal Vector. The figures above demonstrated that the CV curve perfectly aligned with the ground-truth temporal delays imposed between signals. During periods of constant phase delay, the CV remained entirely non-intermittent and parallel to the X-axis, correctly identifying robust temporal stability.

Furthermore, when the simulated delay was subjected to abrupt step changes and gradual phase drifts halfway through the sequence, the algorithm dynamically tracked these modifications with only little lag. Quantitative assessment of the simulation yielded a negligible Mean Absolute Error (MAE) and Root Mean Squared Error (RMSE) between the estimated CV and the true mathematical delay; the values were gathered in Table I (for raw signals and linear modelling; the same could be estimated for phases as well as signal distances).

**TABLE I.** The error between estimated cv and true mathematical delay in synthetic signals (for raw signals and linear modelling).

| Condition | MAE | RMSE |
| --- | --- | --- |
| Without noise, T1 | 0.010 | 0.010 |
| Without noise, T2 | 0.010 | 0.010 |
| Without noise, T3 | 0.010 | 0.010 |
| With noise, T1 | 0.055 | 0.065 |
| With noise, T2 | 0.052 | 0.061 |
| With noise, T3 | 0.056 | 0.068 |

The equality of MAE and RMSE for signals without noise should be interpreted that the algorithm is maximally stable and maintains constant precision.

### B. Physiological Results

The *tempord* framework was applied to the preprocessed (as described in Section 2.4) in-vivo datasets to evaluate its performance and applicability in tracking non-stationary physiological signals. The signals were demonstrated in Figure 5.

**Fig. 5.**
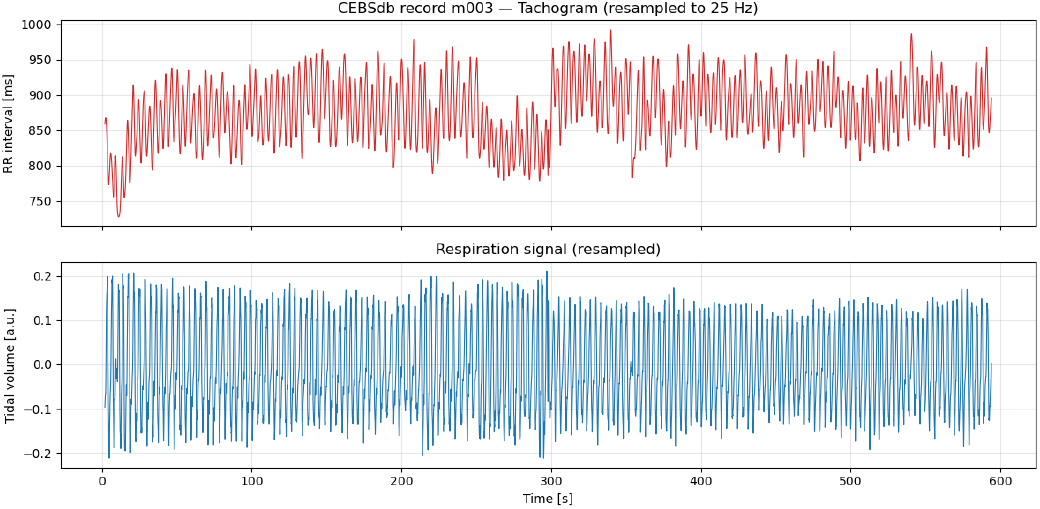
Signals utilized, spontaneous breathing for the first 5 minutes, relaxation for the second 5 minutes (b003+m003 from the CEBSdb).

Applying the algorithm to the sample CEBSdb dataset revealed temporal stability within the cardiorespiratory data during both phases with an optimal shift around half a second. The analyses were performed for raw signals and for linear modeling, for larger threshold than previously used at the level of 0.75 and presented in Figure 6. The extracted Causal Vector exhibited natural, relatively minor fluctuations consistent with standard autonomic variability, as well as data analysis impact.

**Fig. 6.**
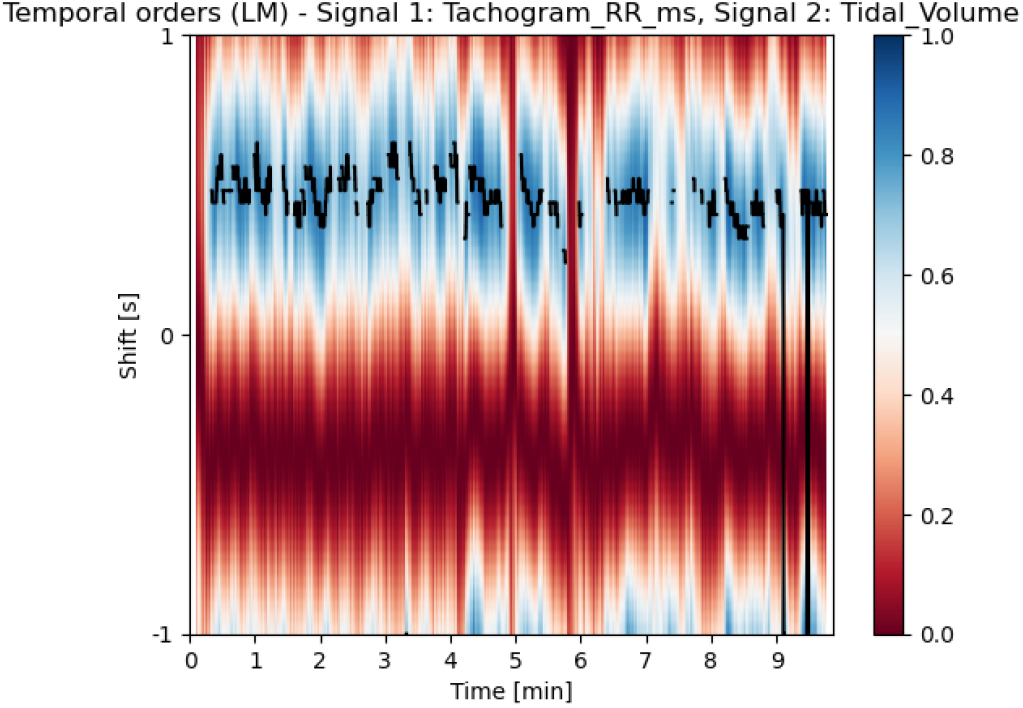
Bivariate temporal order heatmap and CV trajectory for b003+m003 (CEBSdb) signals, illustrating the minor step change in optimal shift during the transition from spontaneous to music-induced breathing after the 5th minute, for linear modeling approach.

Although at first glance the raw signals look quite similar, the mean and standard deviation of the CV for both phases are:

- 0.485 *±* 0.085 s (52.1% above threshold)
- 0.411 *±* 0.161 s (43.5% above threshold), respectively.

At the very end, CV transitions to the second period are also visible (the period decreased in the second phase). This effect would not be present in the result once the TD approach is selected. In Figure 7, the heatmap with “cosine” type, threshold at the level of 0.5, and phase shifts in range of - 2 to 2 seconds, were shown illustratively.

**Fig. 7.**
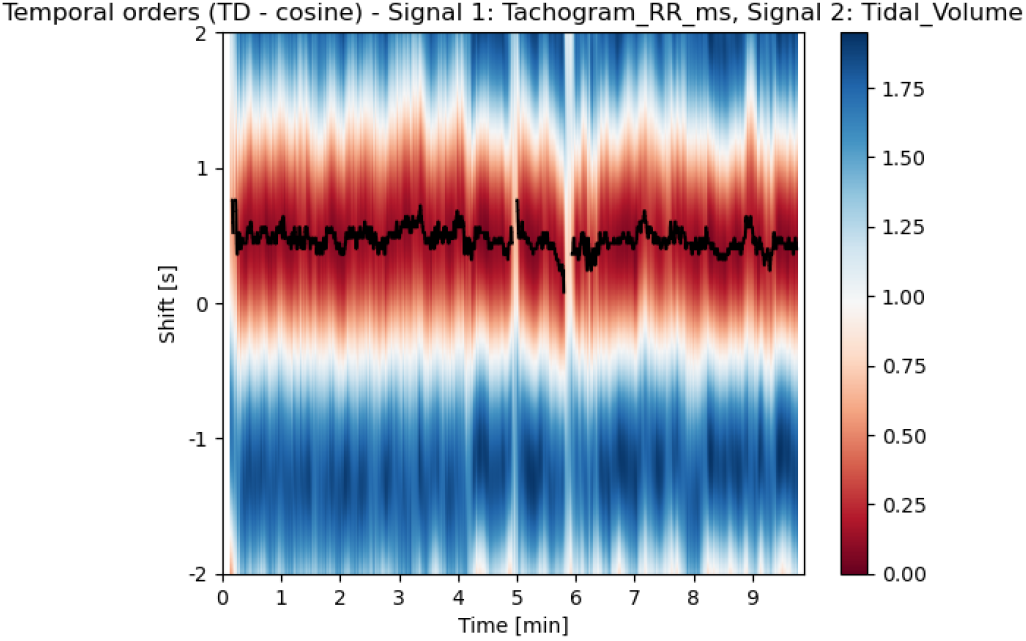
Bivariate temporal order heatmap and CV trajectory for b003+m003 (CEBSdb) signals for time series “cosine” distance approach.

Results of exploratory analysis of computation time in relation to signal length, maximal shifts and sampling frequency are presented in Figure 8. Function execution time varied from 138.5 to 145.8 seconds for signal length, increased from 139.3 to 266.7 seconds and from 6.1 to 138.7 seconds in case of maximal shift and sampling frequency respectively.

**Fig. 8.**
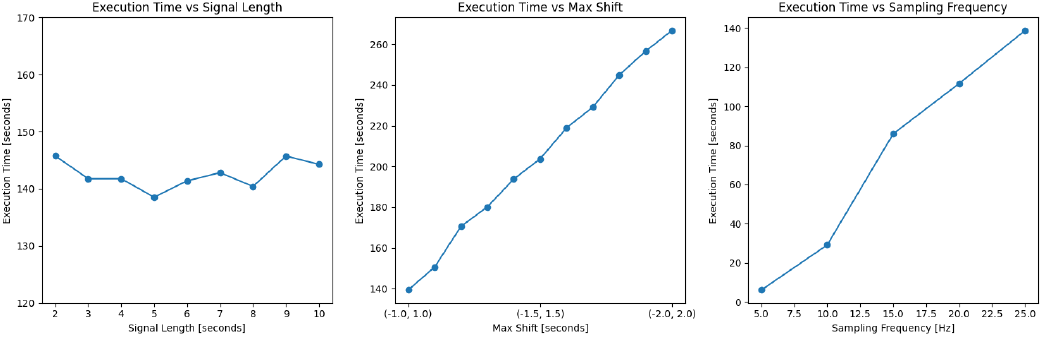
Average computation time across three iterations as a function of signal length (left), maximum shift (middle), and sampling frequency (right).

## IV. Discussion

The introduction of the *tempord* package in Python significantly expands the classical visual and analytical toolkit used in the field of Network Physiology [15]. So far, the methods for exploring temporal relationships within time series, such as linear Granger tests, nonlinear one based on neural network kernel [8] or the advanced Temporal Causal Discovery Framework (TCDF) [3], still focusing primarily on a global perspective. They attempt to reconstruct network structures or averaged dependencies for entire signal segments being analyzed. Such approaches represent a “top-down” strategy, which, while effective in general network identification, often masks the dynamic, short-term fluctuations of biological signals. The local approach proposed in this paper serves as a complementary tool, implementing a “bottom-up” strategy. Instead of imposing rigid, time-invariant structural assumptions, it builds an understanding of dynamic relationships directly from the smallest reasonable time intervals.

The premise underlying *tempord*, that the temporal order of a causal link is a local rather than a global property, is supported by evidence that cardiorespiratory interactions fluctuate even within recordings acquired under nominally constant conditions. Orini et al. [16] showed that the coupling between RR interval and systolic arterial pressure variability is characterized by a time-varying phase difference and time delay, the latter reversing sign in the high-frequency band during head-up tilt and concluded that non-stationary techniques are warranted even for signals conventionally assumed to be stationary (the very assumption embedded in a single, globally estimated model order). The same group extended this to time-frequency coherence between RR variability, arterial pressure and respiration, with automatic significance testing of local estimates [17]. Keissar et al. [18] applied wavelet transform coherence to heart rate and respiration explicitly to recover the transient linear order of the regulatory mechanisms, although their formulation remains in the coherence domain and yields no explicit, interpretable lag.

A key aspect of local analysis using the defined Causal Vector is the ability to precisely evaluate the short-term stability of couplings. Assessing complexity in physiological systems through biomedical signals analysis, particularly during the analysis of CRC, relationships between systems are rarely static [19]. The strength and directionality of these couplings change smoothly or abruptly under the influence of, e.g., body position, physical exertion, study conditions, or specific sleep stages. Global averaging windows can obscure these transient phenomena. Step changes in the CV trajectory (or even fragments where it does not exceed the threshold specified by the user) may directly reflect changes in study conditions, providing researchers with unique behavioral and mechanistic indicators to analyze such part deeper. The approach could hence be utilized to parameterize interventions like head-up tilt test or vagal blockade.

Contrast between spontaneous breathing and breathing during exposure to classical music employs a well-characterized awake paradigm. Bernardi et al. [20] recorded ventilation, RR interval, arterial pressure and baroreflex sensitivity simultaneously during musical excerpts of differing tempo, finding arousal with faster tempi and relaxation during slow passages and pauses; a follow-up study showed that cardiorespiratory variables track the instantaneous music profile, with phrases near the Mayer-wave period entraining autonomic variables [21]. The relevant time scale is therefore that of a musical phrase, not of the whole session, and a session-averaged index necessarily blurs the transitions we aim to resolve. A comparable dissociation appears under cognitive load: Widjaja et al. [22] found that mental arithmetic altered the predictability of the RR series conditioned on respiration without changing coupling strength itself, indicating that strength and temporal structure are not interchangeable descriptors, and that a measure sensitive to the latter may separate conditions the former cannot.

Where the temporal order itself is the quantity of interest, information-theoretic frameworks provide the closest comparison. Faes et al. [23] decomposed transfer entropy into lag-specific contributions across heart period, systolic pressure and respiration during tilt, tracking how the latency of baroreflex and respiratory sinus arrhythmia mechanisms changes with orthostatic stress; earlier work addressed non-linear Granger causality in short cardiovascular series without over-constraining the embedding [24]. Studies of Porta et al. are particularly relevant to the design adopted here. First, omitting respiration biases heart-period–pressure causality, inflating apparently closed-loop interactions during both spontaneous and paced breathing [25]; second, both the choice of coupling estimator [26] and the model-identification strategy [27] affect the resulting causality markers, so any claim about a local link must be reported together with the procedure that produced it.

The transition of the codebase from R to Python represents a strategic leap forward in the package’s utility. In recent years, the data engineering and ML communities have seen a clear shift in preference toward the Python ecosystem, as evidenced by the rapid development of libraries like causal-learn [28] and neurokit2 [14]. The most significant advantage of this architectural change is the ability to directly plug local temporal orders and CV parameters into modern ML pipelines. The locally estimated features can serve as advanced input descriptors for classification models, for instance, in assessment of athletic adaptation.

It should be emphasized that the current version of the framework focuses on bivariate analysis, examining relationships directly between pairs of signals, e.g., tidal volume and RR intervals. This poses a certain limitation in the context of a complete analysis of complex physiological systems, where multiple interdependent feedback loops occur simultaneously. To overcome this barrier, one can evaluate such multivariate relationships within one-vs-one paradigm. Moreover, the parameters obtained from temporal orders analysis could also be aggregated to “global” values by taking a mean of CV values and optimal shift between signals.

The other limitation is a moderate dependence of the results on initial settings, such as the analysis window size (segment length) and the maximum shift range. Selecting too wide a shift range can reveal the influence of subsequent periodic cycles (e.g., subsequent respiratory phases), which significantly complicates the graphical interpretation of heatmaps and distorts the trajectory of the causal vector. Narrowing the analysis window trades statistical power for temporal resolution; the short-series estimators of Faes et al. [24] and the estimator comparisons of Porta et al. [26] both indicate that the variance of causality markers grows quickly as the segment shortens. Our choice of 10-second window length reflects this compromise. Therefore, the effective use of the tool requires researchers to possess preliminary expert knowledge regarding the nature of the studied biological processes to correctly select the shift ranges. Still, the values suggested in the Jupyter Notebook can be considered as ones “to start with” (at least for cardiorespiratory analysis).

Furthermore, the high sampling resolution of modern medical signals poses challenges for the computational efficiency of the algorithm, especially when larger maximum shifts are applied. Therefore, one area of further framework development should focus on targeted code optimization for dense datasets, including the parallelization of computations.

## V. Conclusions

The *tempord* Python package provides a robust, open-source framework for evaluating local temporal orders and causal vectors in bivariate time series, including physiological signals. It is readily accessible to researchers via the Python Package Index (PyPI) and is accompanied by interactive Jupyter Notebooks containing practical code examples to facilitate seamless integration into data science pipelines.

The comprehensive simulation study confirmed the algorithm’s fundamental methodological reliability, demonstrating its precision in tracking mathematically imposed, dynamic phase delays. Furthermore, its application to real in-vivo data highlighted its significant potential utility in exploring dynamic, non-stationary physiological phenomena.

By effectively parameterizing short-term temporal stability and detecting local step changes and CV trends, *tempord* can bridge the gap between global causal discovery and beat-by-beat statistical analysis, offering a possible valuable “bottom-up” instrument for investigating the transient mechanisms governing complex biological networks.

## Declarations

### Data Availability Statement

This study utilizes exclusively simulated data and fully anonymized, publicly available datasets (e.g., PhysioNet). The code, including an interactive Jupyter Notebook demonstrating the entire analytical workflow, is publicly available.

### Ethics Statement

Ethical review and approval were not required for this study, as it does not involve any newly collected human or animal data, relying solely on synthetic and publicly accessible, anonymized datasets.

